# The Nuclear Pore Complex consists of two independent scaffolds

**DOI:** 10.1101/2020.11.13.381947

**Authors:** Saroj G. Regmi, Hangnoh Lee, Ross Kaufhold, Boris Fichtman, Shane Chen, Vasilisa Aksenova, Elizabeth Turcotte, Amnon Harel, Alexei Arnaoutov, Mary Dasso

## Abstract

Macromolecular transport between the nucleus and cytoplasm is mediated through Nuclear Pore Complexes (NPCs), which are built from multiple copies of roughly 34 distinct proteins, called nucleoporins^1–3^. Models of the NPC depict it as a composite of several sub-domains that have been named the outer rings, inner ring, cytoplasmic fibrils and nuclear basket. While the NPC has been extensively studied, the roles of individual nucleoporins within NPCs and their functional interactions remain poorly understood. Here, we applied a rapid degron system to systematically investigate how individual nucleoporins contribute toward NPC architecture. We find that acute depletion of outer ring components (NUP96 or NUP107) disperses the outer ring and cytoplasmic fibrils without disassembly of inner ring members. Conversely, rapid degradation of the inner ring complex component NUP188 disrupts the inner ring without dislodging outer ring members. We also found that depletion of NUP93 destabilized all NPC domains, indicating that it has a unique role as a lynchpin of NPC structure. Our data highlight the modular nature of NPC organization, suggesting that the outer and inner ring complexes do not extensively rely on each other for structural stability after NPC assembly is complete. This dynamic assessment provides new insights regarding the remarkable structural independence of domains within the NPC.

---

Mammalian NPCs are massive structures that span the nuclear envelope, providing not only conduits for transport between the nucleus and the cytoplasm but also an important platform for transport-independent functions in genome organization, gene expression and signal transduction pathways^1,3-8^. Models represent the NPC as a series of concentrically stacked structures aligned around the central transport channel. These structural domains include the cytoplasmic fibrils, outer rings, inner ring, channel and nuclear basket ^1,9^. The Y-complex is a NPC sub-complex that forms the outer ring domains of the NPC. The vertebrate Y-complex contains nine core nucleoporins (SEH1, SEC13, NUP37, NUP85, NUP96, NUP107, NUP133 and NUP160), with a tenth subunit (ELYS) required for chromatin recruitment^3,10,11^. The organization of the Y-complex has been well studied, allowing detailed modeling of this complex within NPCs of many organisms ^1,5,7,12–14^. Structural models typically represent the outer rings as scaffolds formed from multiple copies of the Y-complex arranged into an interlocking lattice. By contrast, models for how the inner ring complex (NUP205, NUP188, NUP155, NUP93, NUP35) contributes to NPC architecture are less advanced, as is our understanding of structural interactions between NPC domains ^3,4,6,9^. Moreover, NPCs undergo disassembly-reassembly cycle during mitotic division, and a lack of tools for acute manipulation of individual nucleoporins has therefore precluded study of their roles in maintaining structures within pre-existing pores.

To address these questions, we utilized CRISPR/Cas9 to insert a genomic cassette encoding a fluorescent NeonGreen moiety and an Auxin-Inducible Degron (AID) tag homozygously at nucleoporin gene loci in human colorectal adenocarcinoma DLD-1 cells (Extended Data Fig. 1), thereby achieving complete tagging of individual nucleoporins at their N- or C-terminus. We introduced these tags into genes encoding evolutionarily conserved members of the Y-complex ^15^ (NUP160, NUP133, NUP96, NUP107 and NUP85; Fig. 1a, b; Extended Data Fig. 1a). We also used CRISPR/Cas9 to program the cells for TIR-1 protein expression under the control of the ubiquitously expressed RCC1 gene ^16^. TIR-1 is necessary for specific, auxin-dependent degradation of AID-tagged proteins ^17^,^18^. All cells with individually tagged Y-complex nucleoporins showed NeonGreen localization at the nuclear rim during interphase and at kinetochores during mitosis (Fig. 1c; Extended Data Fig. 1b, c)^19–21^. Auxin addition resulted in a rapid loss of the fluorescent protein (Fig. 1c). Degradation after auxin addition was confirmed by Western blotting with antibodies that recognize antigens at the terminus of the nucleoporin opposite from the degron tag (Extended Data Fig. 1e). Both immunoblotting and live microscopy indicated substantial depletion within one hour of auxin treatment and near-complete degradation within 3-4 h (Fig 1d; Extended Data Fig. 1f).

**Figure 1.**
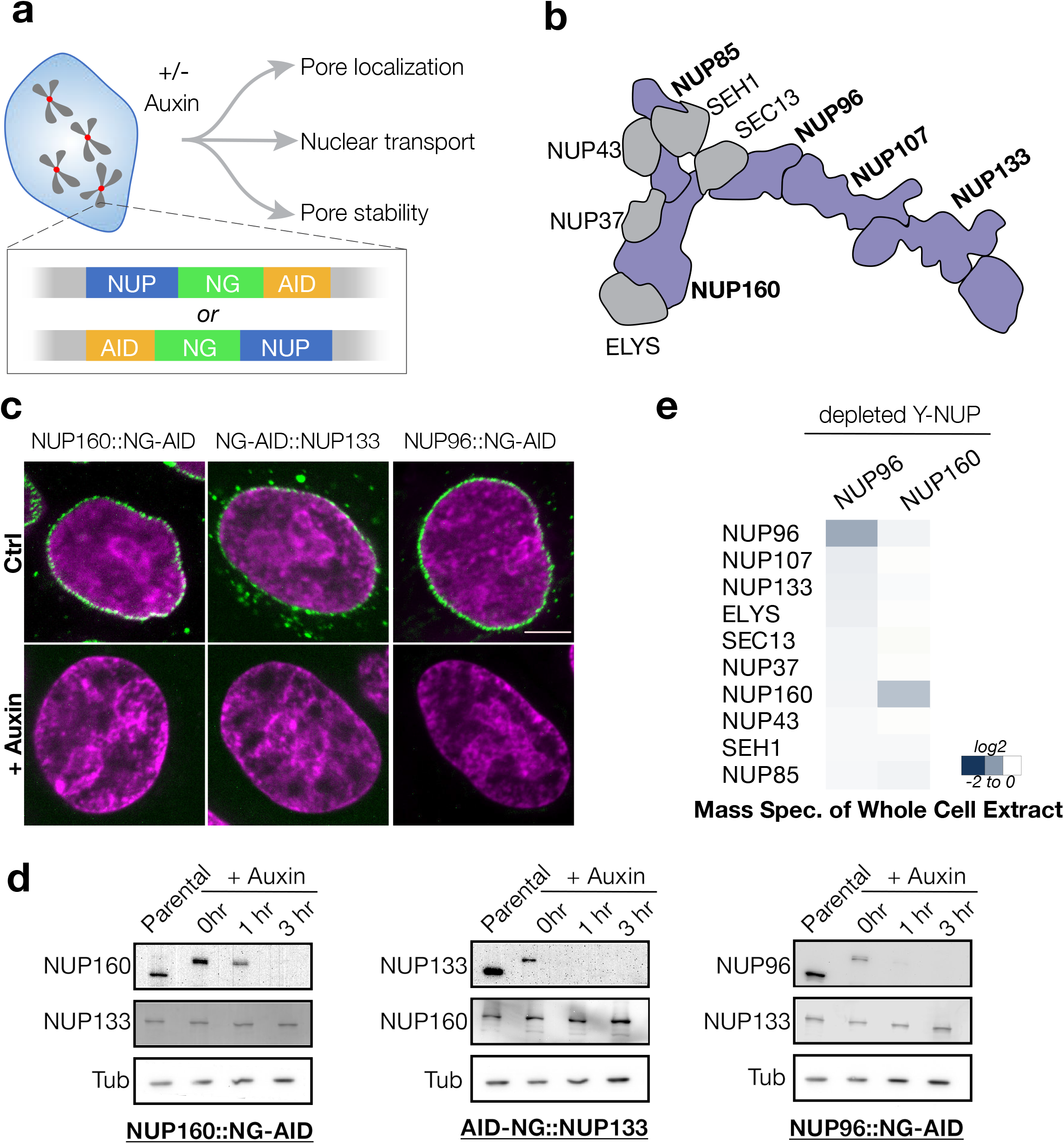
Selective analysis of individual Y-complex nucleoporins. a. Experimental workflow. b. Schematic representation of Y-complex organization (based on von Appen et al.^8^); highlighted in blue are Y-NUPs for which biallelically tagged-cell lines were successfully generated and validated. c. Tagged NUP160, NUP133 and NUP96 nucleoporins (green) with and without Auxin treatment (4 hours, 1mM). Regulator of Chromosome Condensation 1 protein (RCC1) tagged with Infra-red Fluorescent Protein (IFP), RCC1::IFP, serves as a chromatin marker and is shown in magenta ^16^. (Scale bar = 5 μm.) d. Western blotting of protein degradation upon Auxin treatment. Hours of auxin treatment are indicated on top. β-tubulin (Tub,loading control) and another Y-complex member was used to show specific degradation of the indicated Y-NUP. e. A heat map of all Y-NUPs of TMT-based mass-spectrometry of whole cell lysate upon 4 hours of auxin depletion. Specific degradation of Y-NUP was achieved with minimal effect to other Y-members.

In previous RNAi-based experiments, depletion of individual Y-complex members required days of incubation and caused concurrent loss of other nucleoporins in this complex^22,23^. We checked whether the same phenomenon would occur during the much more rapid time course of depletion after auxin addition to cells with AID-tagged nucleoporins. Western blotting indicated no substantial alterations in the protein levels of non-targeted complex members after auxin addition (Fig. 1d). We used Tandem Mass Tag (TMT)-assisted mass spectrometry of total cell lysates to extend this observation: we detected substantial depletion only for the AID-tagged complex member while other Y-nucleoporins remained relatively unchanged (Fig. 1e). Therefore, the AID-based tagging permits fast, efficient and highly selective degradation of individual nucleoporins. Long-term depletion of any individual Y-complex member led to growth arrest of cells with an overall loss of cell viability, recapitulating prior RNAi-based results regarding the phenotype after extended depletions ^23^ (Extended Data Fig. 2a-e). In most cases, this growth arrest appeared to coincide with a failure to reassemble NPCs and expand nuclear volume in post-mitotic auxin-treated cells (Extended Data Fig. 2f, g). The pronounced defects in post-mitotic NPC assembly indicate that while nucleoporins share evolutionarily conserved features ^15^, they do not act redundantly during de novo NPC formation.

In viewing Y-complex structural models and proposed mechanisms for its assembly into NPC outer rings ^8,14,24^, we anticipated that depletion of any Y-complex member should result in complete destabilization of the outer rings. To our surprise, depletion of NUP160, NUP133 or NUP85 did not cause obvious changes in the localization of other Y-members at the NPC (Fig. 2a; Extended Data Fig. 3). By contrast, depletion of NUP96 dispersed NUP133 from the NPC, as we had originally anticipated (Fig. 2a). To validate these observations, we used CRISPR/Cas9 to generate dual-tagged constructs wherein a second nucleoporin is fluorescently labelled. Consistent with immunostaining results, the NUP96::BFP signal persisted at the nuclear rim upon depletion of NUP160 (Fig. 2b), but loss of NUP96 dispersed mCh::NUP133 (Fig. 2b). These findings suggest that the outer ring structure is remarkably resistant to perturbations once it is fully assembled, and show that its members are not of equivalent importance in sustaining its stability.

**Figure 2.**
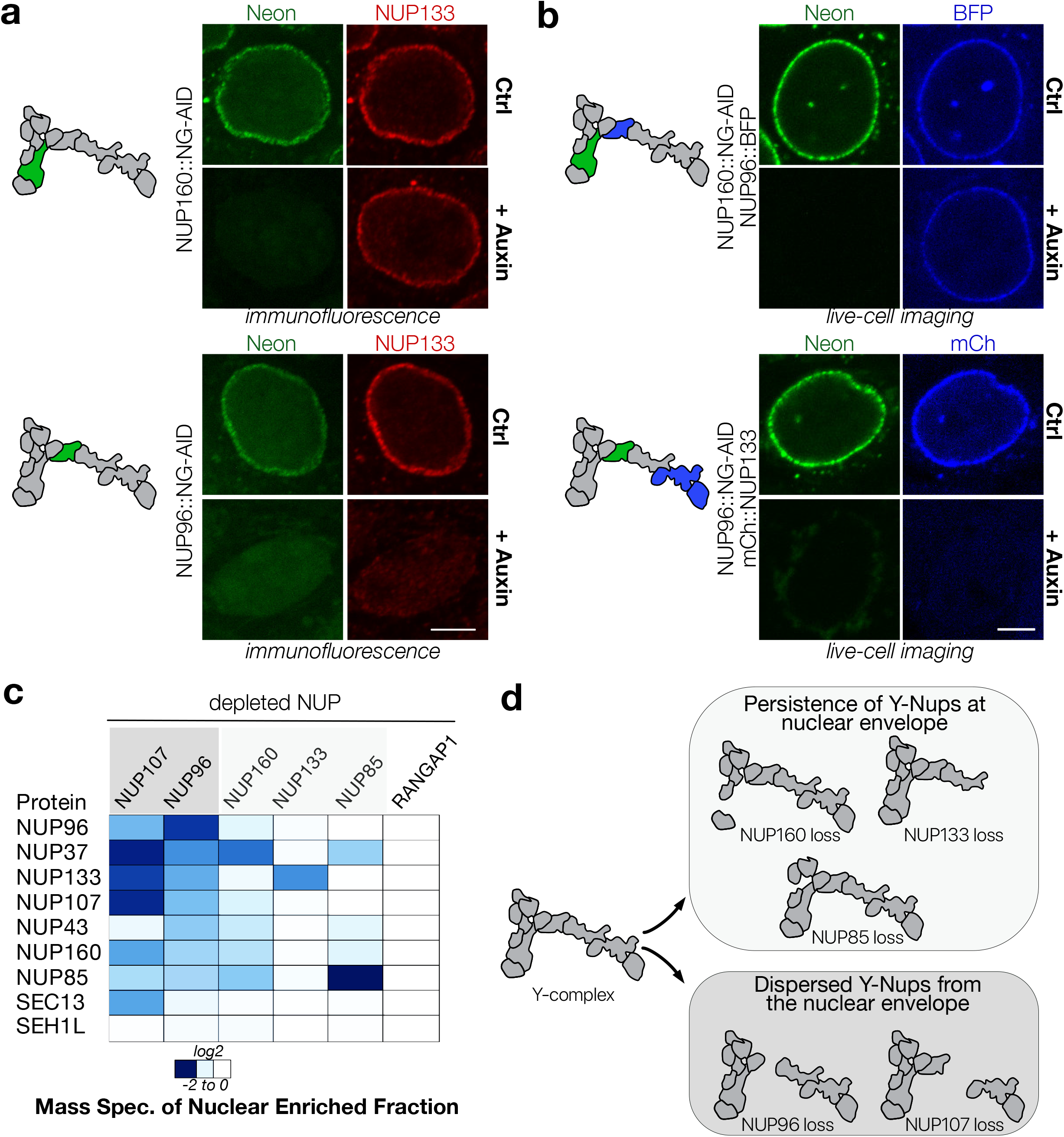
Stability of nuclear pore upon rapid loss of Y-NUPs. a. Immunofluorescent localization of NUP133 (red) in NUP160::NG-AID and NUP96::NG-AID (green) cell lines upon auxin treatment (fixed cells, 4hr, 1mM). Localization of NUP160 and NUP96 depicted in the schematic, respectively. Scale bar: 5 μM. b. Localization of NUP96 upon loss of NUP160 (top) and localization of NUP133 upon loss of NUP96 (bottom) in live-cell experiments using auxin (4hr, 1mM). Scale bar: 5 μM. BFP (Blue Fluorescent Protein), mCh (mCHERRY). c. A heat map of all Y-NUPs of TMT-based mass-spectrometry of nuclear-pore enriched fraction upon auxin depletion of the tagged 5 Y-NUPs. d. Schematic depicting differential effects on the Y-NUP overall upon loss of individual Y-members.

To obtain a quantitative and comprehensive assessment of how the Y-complex is re-distributed upon loss of individual members, we purified NPC-enriched fractions from control and auxin-treated cells and performed TMT-labelling followed by mass spectrometry. In agreement with live-imaging and immunofluorescence experiments, the strongest change in the Y-complex occurred after NUP96 or NUP107 depletion (Fig. 2c). There was also a marked alteration in the abundance of Y-complex members within the NPC-enriched fractions after NUP160 depletion, but only a mild effect with NUP133 and NUP85 depletion. Together, these results indicate that a subset of Y-complex members, such as NUP96 and NUP107, are essential for the persistence of the outer ring lattice. However, NUP133 and NUP85 appear to be less critical for sustaining the lattice once it is in place (Fig. 2d), with NUP160 demonstrating an intermediate phenotype.

Importantly, near-complete loss of the outer ring in NUP96-depleted cells did not cause collapse of the whole NPC, as demonstrated by persistent immunostaining with mAb414 antibodies to detect nucleoporins with FG repeats (NUP62, NUP153, NUP214 and RanBP2)^25^, indicating remarkable resilience of the NPC structure (Fig. 3a). To quantify global changes after outer ring loss, we further analyzed TMT-mass spectrometry results to assess nucleoporins in other NPC domains (Fig. 3b). Degradation of NUP96 and disruption of the outer rings caused a loss of cytoplasmic fibril nucleoporins (RANBP2, RANGAP1, NUP88, NUP214) and a ~50% reduction of basket nucleoporins (TPR, NUP153, NUP50). However, the effect of loss of NUP96 on both the inner ring and channel nucleoporins was minimal.

**Figure 3.**
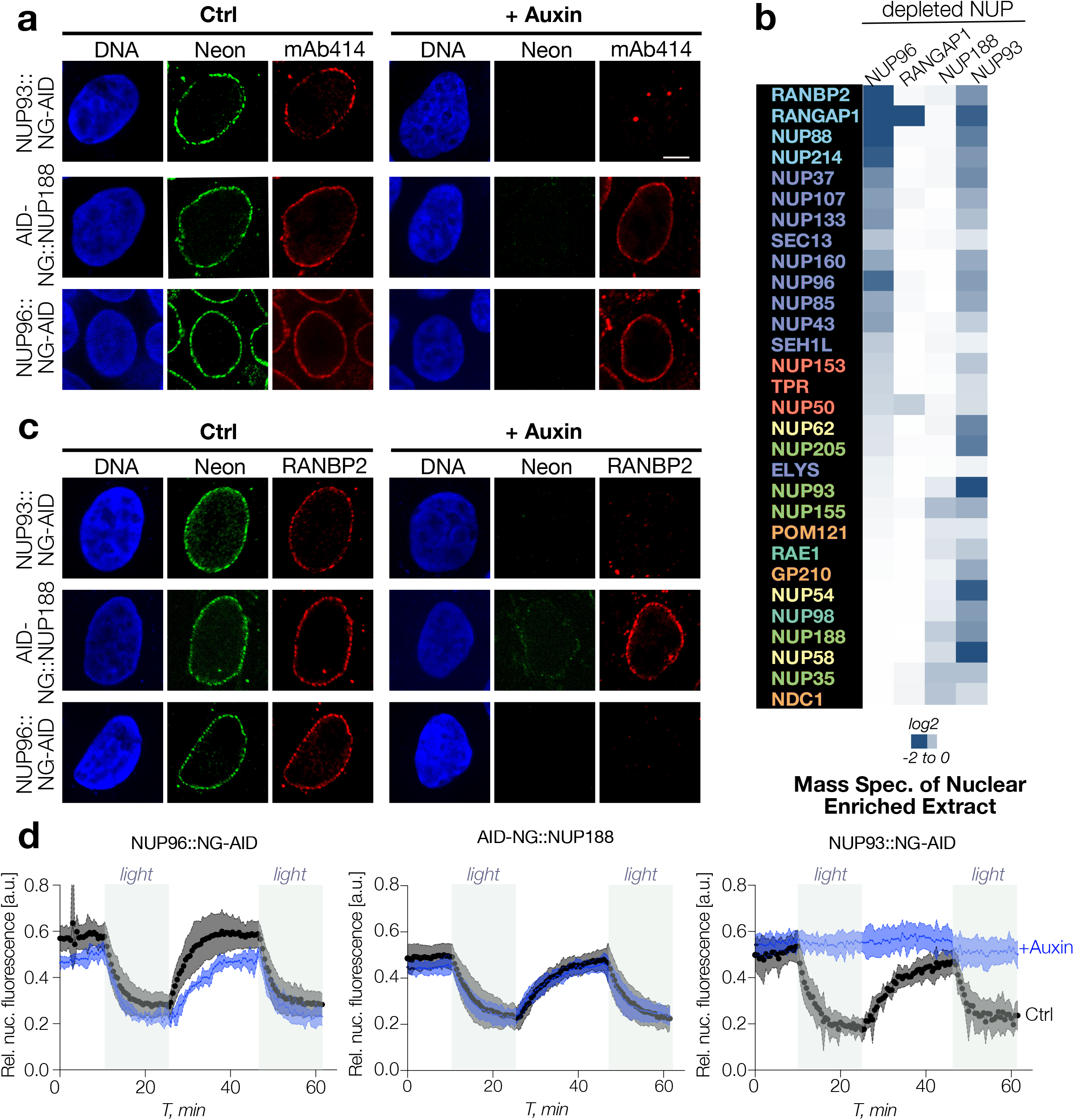
Effects of the loss of NUP96, NUP93 and NUP188 on the nuclear pore architecture. a. mAb414 signal upon depletion of NUP93, NUP188 and NUP96 respectively) upon auxin treatment (6 hr, 1mM). Scale bar: 5 μM. b. A heat map of all nucleoporins of TMT-based mass-spectrometry of nuclear-pore enriched fraction upon auxin depletion of NUP96, NUP188 and NUP93. RANGAP1 had minimal effects on other nucleoporin-profile and was used as a negative control. c. Localization of RANBP2 upon depletion of NUP93, NUP188 and NUP96 respectively) upon auxin treatment (6 hr, 1mM). Scale bar: 5 μM. d. Nuclear transport of a model substrate^26^ in the presence and absence of NUP96, NUP188 and NUP93, respectively. Y-axis denotes timing and X-axis denotes relative nuclear fluorescence (arbitrary units).

The persistence of the inner ring after outer ring disruption led us to test the autonomy of these structures further using AID degron cell lines for inner ring nucleoporins NUP188 and NUP93 (Extended Data Fig. 4a, b). NUP188 depletion caused a NPC disassembly profile that was opposite to the profile after NUP96 depletion: inner ring components were extensively displaced, while the components of the cytosolic fibrils, outer ring and basket were largely unaffected (Fig. 3b). Interestingly, there was a global reduction of almost all nucleoporins upon loss of NUP93. Consistent with the TMT mass spec profiles, staining with mAb414 antibodies showed almost complete loss of NPC signal after NUP93 depletion, but this was not observed after NUP96 or NUP188 depletion (Fig. 3a). RanBP2, a cytoplasmic filament nucleoporin, was lost upon depletion of either NUP96 or NUP93 (Fig. 3b), consistent with earlier structural predictions and it was also sensitive to depletion of NUP107 (Extended Data Fig. 3b). Together, our results demonstrate that the inner- and outer rings of the NPC form distinct and independent structures, and NUP93 serves as a NPC lynchpin that is essential for both of them.

We tested whether the residual structures after depletion of the inner ring or outer rings remained functional for the import and export of a model fluorescent substrate whose distribution is regulated by exposure to blue light ^26^. Remarkably, there was only a minimal change in both nuclear import and export rates upon loss of NUP96, and essentially no change in nuclear transport for the model substrate after NUP188 depletion (Fig. 3d). However, NUP93 depletion caused a complete block in nuclear transport in both directions, confirming that this global disruption functionally disabled NPCs. Together, these results indicate that persistent inner ring or outer ring structures could still act as conduits for vectoral nuclear trafficking and that these modules can support independent and redundant trafficking routes. However, removal of both sets of structures forecloses all nuclear trafficking.

To evaluate structural alterations at the pore after depletion of NUP96 and NUP93, we performed high resolution scanning electron microscopy (SEM). Depletion of NUP96 caused a minor reduction (~30%) in the overall number of NPCs, which might reflect inhibition of de novo pore assembly during auxin treatment. More interestingly, there were significant changes in the architecture of remaining NPCs, which showed decreased size and loss of cytoplasmic fibrils (Fig. 4a, b, Extended Data Fig. 6b). By contrast, there was a precipitous drop (> 8-fold) in the number of NPC-like structures upon NUP93 depletion (Fig. 4d); these sparsely distributed NPC remnants displayed narrowing to various extents, which might suggest that they were becoming sealed with the membrane (Fig. 4c, Extended Data Fig. 6c). These images provide a dramatic visual confirmation of NPC disruption after the loss of individual nucleoporins that comports well with biochemical data from mass spectrometry (Fig. 3b) and functional data from nuclear trafficking (Fig. 3d).

**Figure 4.**
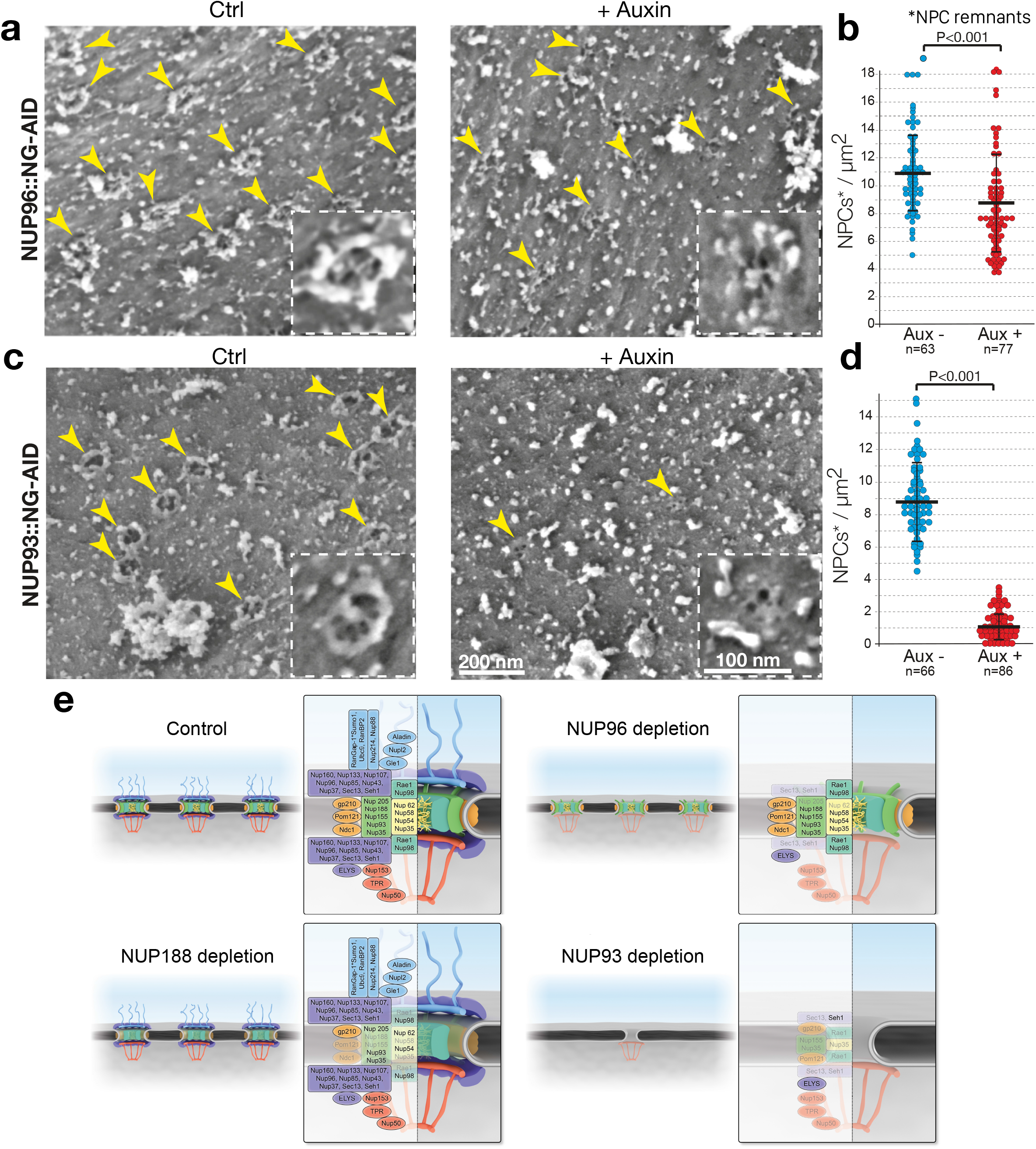
Effects of the loss of NUP96 or NUP93 on NPC architecture. Direct surface imaging of exposed nuclei by SEM was performed on control (left pane) and auxin treated NUP96::NG-AID (a) and NUP93::NG-AID (b) cell lines. Auxin was treated to ensure complete degration (3hr for NUP96 and 6hr for NUP93). Yellow arrows depict individual NPCs or NPC-like remnants. Insets, bordered by dotted white lines show higher magnifications of individual NPCs or NPC remnants. Scale bars as indicated. (c) and (d) depict the quantification of NPCs/μm^2^ in control and auxin treated conditions using a two-tailed Student’s t-test. n = number of nuclear envelope fields analyzed in three independent experiments. See Supplementary Fig. 5 for additional details. (e) A suggested model for NPC structure upon loss of NUP188, NUP96 or NUP93.

Our findings provide important and unanticipated insights into nuclear pore structure: The outer ring nucleoporins were critical for building a fully functional NPC, yet established NPCs were resilient even after dispersion of the outer rings, or, for the most part, of the inner ring. Two Y-complex nucleoporins, NUP107 and NUP96, acted as critical structural pillars of the outer rings of established NPCs (Fig. 2). Detailed analysis of NPCs after NUP107 or NUP96 depletion indicated that the outer ring reinforces NPC structure, stabilizing the cytoplasmic fibrils and, to a lesser extent, nuclear basket. Conversely, loss of NUP188 extensively dispersed inner ring components, including channel nucleoporins that are involved in nuclear transport, yet left the outer rings intact. Hence, the outer and inner rings appear to represent two different subclasses of nucleoporins, both of which have critical and partially redundant roles in nuclear transport (Fig. 3d). NUP93 held both the inner and outer rings together, such that NUP96 and NUP188-dependent nucleoporins were all extensively dispersed upon NUP93 depletion (Fig. 3b). Moreover, the drastic changes in NPC composition after NUP93 loss were associated with collapse of residual structures and apparent sealing of the membrane, along with a dramatic drop in the number of detectable pores in the nuclear envelope (Fig. 4).

Our finding that NPCs remain extensively functional after depletion of individual nucleoporins (Fig. 3d) contrasts with earlier reports that they are essential for cell viability^22, 23, 27^. This difference can be reconciled through the observation that loss of any inner or outer ring components tested disrupted post-mitotic NPC assembly (Extended Data Fig. 2). Thus, while individual nucleoporins may be indispensable during NPC assembly ^28,29^, NPC maintenance is much more flexible, with the persistence of functional remnants for extended periods of time. This distinction would be difficult to observe in RNAi-based experiments, given the extensive incubation time required for nucleoporin depletion spanning multiple cell cycles ^22,23,27^. Importantly, the reported recruitment timing for a particular nucleoporin during NPC assembly did not correlate well with the extent to which it was essential in established NPCs. For example, Y-complex members are recruited early and synchronously during post-mitotic NPC assembly ^23,30^, leading to the expectation that they should act as a cohesive structure and that loss of any member might cause complete collapse of the complex. This did not prove to be the case: individual Y-complex members could be removed without destruction of the outer ring lattice (Fig. 2) and even the loss of the lattice itself left remarkably functional NPC remnants (Fig. 3). Conversely, NUP93 shows relatively later recruitment during post-mitotic NPC assembly^31^, yet NUP93 acted as a lynchpin whose loss was highly disruptive to all NPC domains (Fig. 3). Assembly and disassembly are thus clearly distinct: NPC structures that do not assemble without prior formation of the outer rings can persist after those rings are removed, and removal of NUP93 leaves remnants that are not equivalent to the assembly intermediates into which it was inserted during NPC formation.

Finally, we note two important implications of our findings: First, the persistence of functional pores lacking a subset of canonical nucleoporins suggests the possibility that terminally differentiated cells might retain substantial nuclear trafficking even with divergent NPC composition, particularly if their turnover of NPCs were slow enough that de novo assembly is not practically limiting ^32^. Differentiated cells might thus customize function through altered NPC composition, potentially modulating specific trafficking pathways or aspects of NPC activity such as gene regulation and post-translational protein modifications ^33–35^. Second, it has long been noted that a handful of structural motifs related to those found in Coat Protein II (COPII) complexes are repeated within the different sub-complexes that form the NPC ^36,37^. It has also been observed that different organisms show strikingly altered organization of these components, for example outer rings that consist of 8 versus 16 copies of the Y-complex ^5,13,38^, and that nucleoporins possess sequence divergence consistent with strong selection pressures ^39^. We observed a capacity of modules within the NPC to function somewhat independently; this independence may reflect the parallel assembly of these modules from related building blocks and may explain how proteins within an essential structure like NPCs can sustain both adaptability and high functional capacity during evolution.

## METHODS

### Cell culture, transfection and immunofluorescence

DLD-1 cells were maintained in DMEM medium with 10% fetal calf serum and Glutamax at 5% CO_2_ and 37°C and as previously described ^16^. Transfection was performed with a slight modification as previously described ^16^. Briefly, 1000 ng of donor and gRNA plasmids in 1:1 ratio was used using ViaFect (Promega) transfection reagent as per manufacturer’s guidelines. Localization and expression of targeted proteins were validated using previously described approaches ^16^.

For immunofluorescence, cells were fixed with 4% paraformaldehyde in PBS with 0.5% Triton X-100 at RT for 15 min. Blocking was done with 10% horse serum for at least 20 min. Nucleoporin-specific antibody was used for primary staining (over-night incubation at 4°C) and and AlexaFluor-conjugated secondary antibodies (Invitrogen; 2 hours at RT). Microscopy was performed as previously described ^16^. Brightness and contrast were applied equally to all images within the experiment using Fiji software. RGB images from Fiji were processed to final resolution using Affinity Designer.

3-Indoleacetic acid powder (Sigma Aldrich) was dissolved in water to a concentration of 500mM and frozen. Fresh aliquots were used to a final concentration of 1mM. 3-4 hrs of 1mM IAA was used to deplete the protein, with the exception of NUP93 for which 6 hrs was required.

### Plasmid construction

CRISPR/Cas9 was used for endogenous targeting. 2 gRNA plasmids were generated per construct and were integrated into pX330 (Addgene# 42230) using BbsI site as previously described ^16^. Guide RNAs were selected using now-defunct CRISPR Design Tools from http://crispr.mit.edu:8079. Universal donor vector (pCassette) was generated by restriction digest of pEGFP-N1 vector (Clontech) to insert a new multiple cloning site as previously described^16^. The sequences of homology arms were amplified using genomic DNA extracted from DLD-1 cells. Three copies of reduced AID tag (3xminiAID) was used^18^. cDNAs of mCherry was amplified from pmCherry-N1 (632523, Clontech). BFP sequence was synthesized using IDT Company’s synthesis service. All PCR reactions were performed using Hi-Fi Taq (Invitrogen) with appropriate primers.

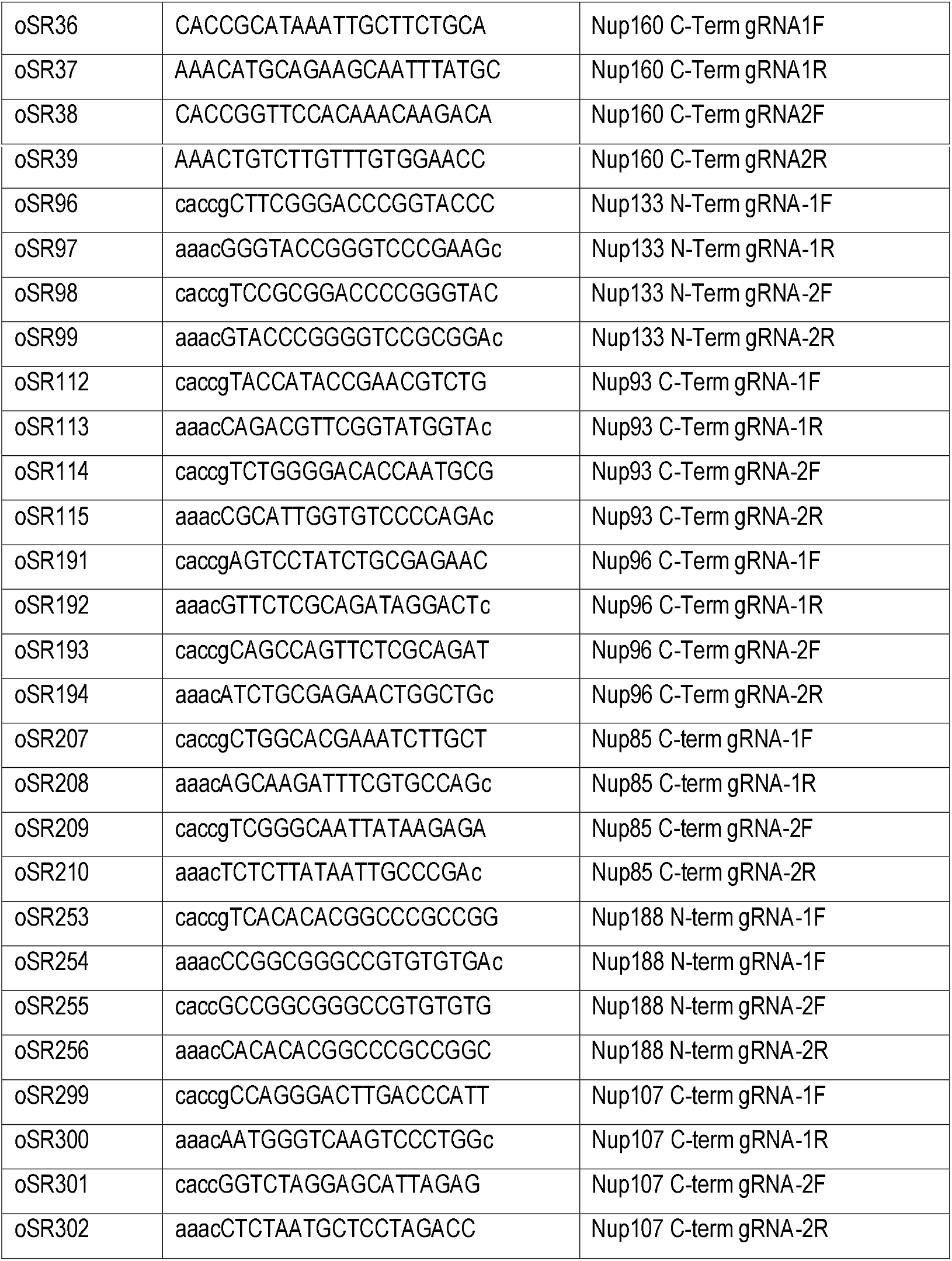

Previously described gRNA pairs used for targeting TPR and RANGAP1 ^16^. TIR1 knock-in was performed as previously described^16^.

### Genotyping

DNA from DLD-1 and nucleoporins-targeted cells was extracted with Wizard^®^ Genomic DNA Purification Kit (Promega). Clones were genotyped by PCR for homozygous insertion of tags with two sets of primers listed below.

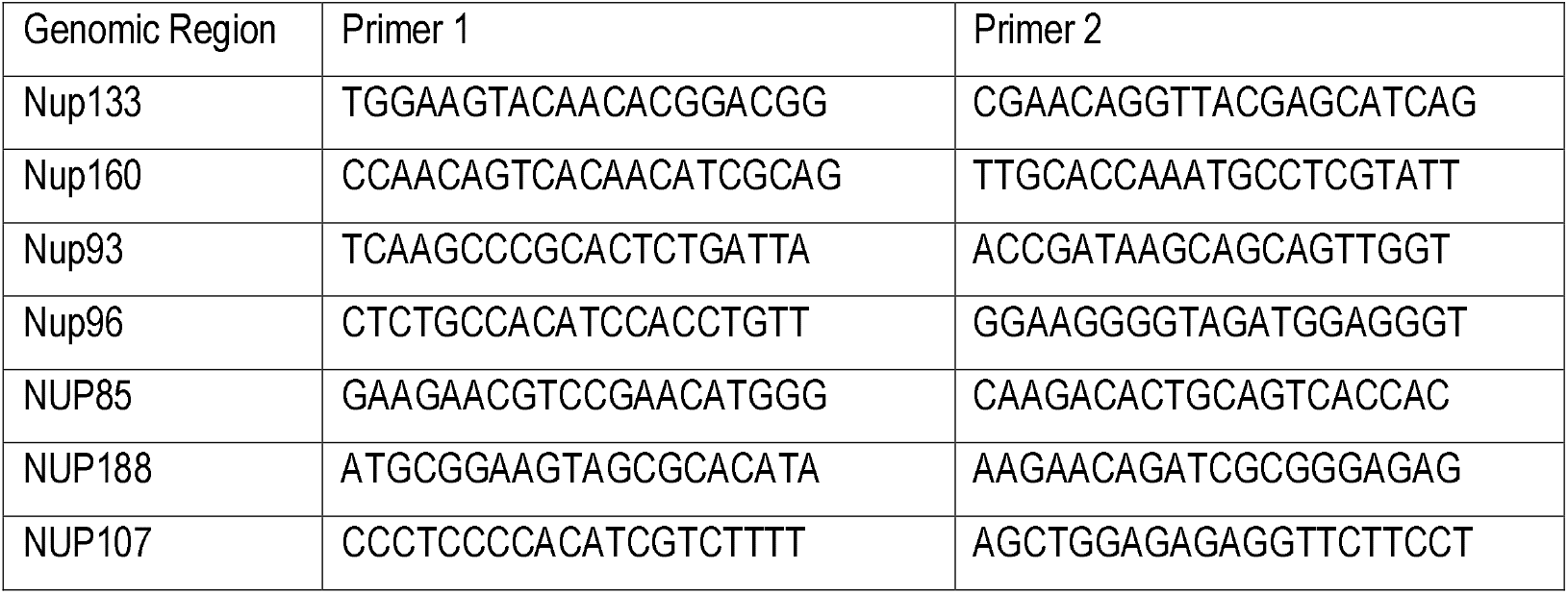

### Crystal violet staining

Crystal violet staining was performed as previously described ^16^.

### Analysis of NPC and NPC-associated proteins by mass spectrometry

Nuclear pore complexes-enriched fraction isolation, mass spectrometry analysis, and protein identification and quantification analysis was done in a previously described manner ^16^.

### Protein extraction and Western blot

2x Laemmli Sample Buffer (Bio-Rad) was used to lyse DLD-1 cell pellets, which were boiled for 15 min at 98°C and ultra-centrifuged at 117K for 15 min at 16°C. SDS-PAGE and Western blotting were performed as described elsewhere ^16^.

### Antibodies

The secondary antibodies for immunofluorescence (AlexaFluor-conjugated) and western-blot analysis (HRP-conjugated) were purchased from Invitrogen and Sigma-Aldrich respectively. Following antibodies were used:

| Protein | Source |
| --- | --- |
| NUP160 | St John's Lab (STJ94576) |
| NUP133 | Bethyl (A302-386A) |
| NUP96 | Bethyl(A301-784A) |
| mAb414 | Biolegend (MMS-120P) |
| Beta-tubulin | Cell Signaling (#2146) |
| RANBP2 | Dasso Lab<sup>40</sup> |

### Scanning Electron Microscopy

DLD-1 nuclei were exposed for direct surface imaging by SEM using previously described methods^41,42^. In brief, cells at 80% confluency were trypsinized and subjected to hypotonic treatment in the absence of detergents and subsequently passed through a 21-gauge needle. The resultant specimens were gently centrifuged onto the surface of 5 x 5 mm^2^ silicon chips, fixed, and further processed for electron microscopy. Dehydration was performed using a graded series of ethanol solutions and critical-point drying was performed using a K850 apparatus (Quorum Technologies). Q150T turbo pumped sputter coater (Quorum Technologies) was used to coat the samples with a ~1 nm thick layer of iridium. Imaging was performed using a Merlin field emission scanning electron microscope (Zeiss) equipped with a secondary electron in-lens detector.

### Statistical Analysis and Reproducibility

Immunofluorescence, SEM and live imaging experiments were performed at least three times.

## Supporting information

Supplementary Figure 1. Y-complex NUPs can be endogenously tagged and rapidly degraded

Supplementary Figure 2. Effects on long-term depletion of individual Y-NUPS

Supplementary Figure 3. Stability of nuclear pore upon rapid loss of Y-NUPs

Supplementary Figure 4. Data supporting NUP93 and NUP188 tagging

Supplementary Figure 5. SEM imaging of an individual nucleus

Supplementary Figure 6. Diversity of structures upon individual nucleoporin depletion

## ACKNOWLEDGEMENTS

We thank all the members of the Dasso lab for comments on the manuscript. We also thank NICHD Microscopy and Imaging Core for its assistance. This work was supported by NICHD project no. HD008954 to M. D.; a research grant from the Israel Science Foundation (958/15) to A. H. and Intramural Research Fellowship (IRF) to S.G.R.

## SUPPLEMENTARY FIGURE LEGENDS

**Supplementary Figure 1. Y-complex NUPs can be endogenously tagged and rapidly degraded.**

a. A schematic showing the protein product resulting from endogenous tagging in DLD cells. With the exception of NUP133, all the other Y-NUPS (NUP160, NUP107, NUP96 and NUP85) were tagged at the C-terminus as indicated in the figure.

b. Live cell imaging of tagged NUP160, NUP133 and NUP96 nucleoporins (green) during interphase (top panel) and mitosis (bottom panel). Chromatin (RCC1::IFP) is shown in magenta. Nuclear rim localization was observed in interphase and kinetochore localization was observed in mitosis (Scale bar: 5 μm).

c. Live cell imaging results depicting localization of NUP85 and NUP107 tagged cell lines in interphase.

d. Image of agarose gel with PCR-products to validate endogenous tagging of Y-NUPs. Specific primer pair was used for each nucleoporin and genomic DNA from wild-type DLD-1 cells was used as control.

e. Schematic of NUP160, NUP133 and NUP96 highlighted with antibody recognition site.

f. Measurement of NUP96::NG-AID degradation in NUP96::NG-AID tagged cells after auxin treatment by live imaging. X-axis represents time (hours) and y-axis represents fluorescence intensity (arbitrary unit).

**Supplementary Figure 2. Effects on long-term depletion of individual Y-NUPS.** Bar graphs indicate cell growth upon days of auxin treatment in control (green) and auxin treated (white) cell lines for NUP96 (a), NUP160 (b), NUP107 (c), NUP133 (d) and NUP85 (e). Kinetics of nuclear growth upon loss of NUP160 (f) and NUP96 (g) using auxin in mitotic cells. 0 minute was used as time of cells in metaphase.

**Supplementary Figure 3. Stability of nuclear pore upon rapid loss of Y-NUPs.**

a. Localization of NUP98 upon depletion of NUP96. NUP96::NG-AID cell lines were generated by C-Terminal tagging, therefore, the effect on NUP98 stability and localization is nominal.

b. Localization of RANBP2 upon depletion of NUP107. Depletion of NUP107 results in loss of nuclear localization of RANBP2 in a similar manner as NUP96 depletion.

c. Localization of NUP133 upon depletion of NUP85. NUP133 signal remains at the pore, in a similar manner as depletion of NUP160. d. Localization of NUP107 upon depletion of NUP133. NUP107 signal remains at the pore. In all the cases, depletion was achieved using auxin treatment for 6 hours, scale bar: 5uM.

**Supplementary Figure 4. Data supporting NUP93 and NUP188 tagging.** Image of agarose gel with PCR-products to validate endogenous tagging of inner nucleoporins (a) NUP188 and (b) NUP93. Specific primer pair was used for each nucleoporin and genomic DNA from wild-type DLD-1 cells was used as control.

**Supplementary Figure 5. SEM imaging of an individual nucleus.**

a. A representative individual exposed nucleus captured at a magnification of 16,660x showing a collection of rectangular fields containing relatively flat nuclear envelope expanses. Multiple fields of this type were used for the quantification of NPCs/μm^2^ in Figure 4. Note that because of steep angles and obscuring debris it is impossible to use the whole visible area of any individual nucleus for such measurements. b. Higher magnification of the rectangular field shown in red, above. The area delineated by a dashed white line was chosen for measurement. All the images used for this analysis were captured at a magnification of 100,000x.

**Supplementary Figure 6. Diversity of structures upon individual nucleoporin depletion.**

Galleries of enlarged, individual NPCs or NPC-like remnants upon depletion of NUP96 and NUP93 pre and post-auxin treatment, selected from the experiments summarized in Figure 4.

## Acronyms

NPC: nuclear pore complex
NG: NeonGreen
AID: Auxin Inducible Degron
TMT: Tandem Mass Tag
TIR1: Transport Inhibitor Response
RCC1: Regulator of Chromosome Condensation 1

