## Supplementary figures and images for "The Nuclear Pore Complex consists of two independent scaffolds"

### Supplementary Figure 1. Y-complex NUPs can be endogenously tagged and rapidly degraded

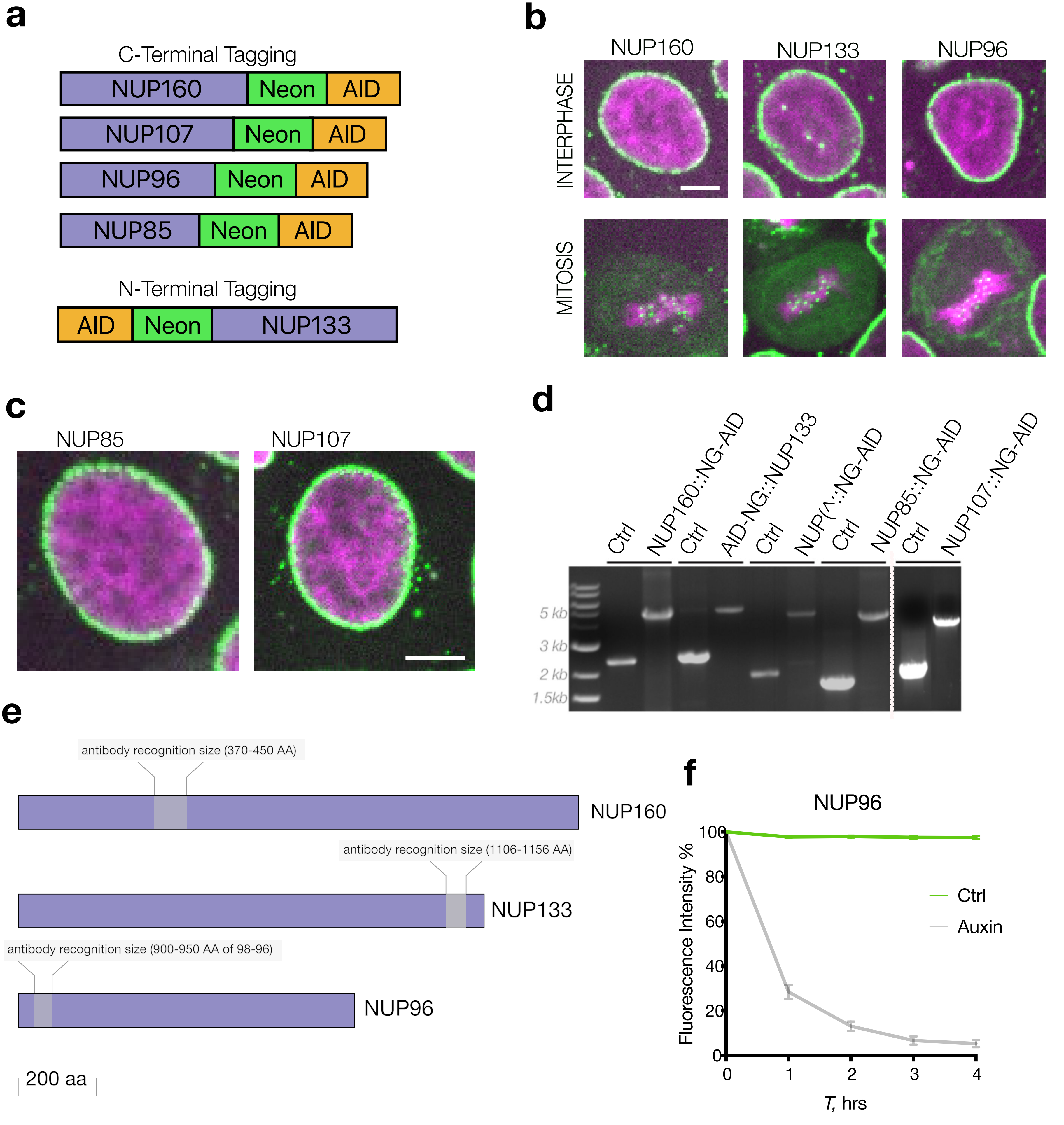

### Supplementary Figure 2. Effects on long-term depletion of individual Y-NUPS

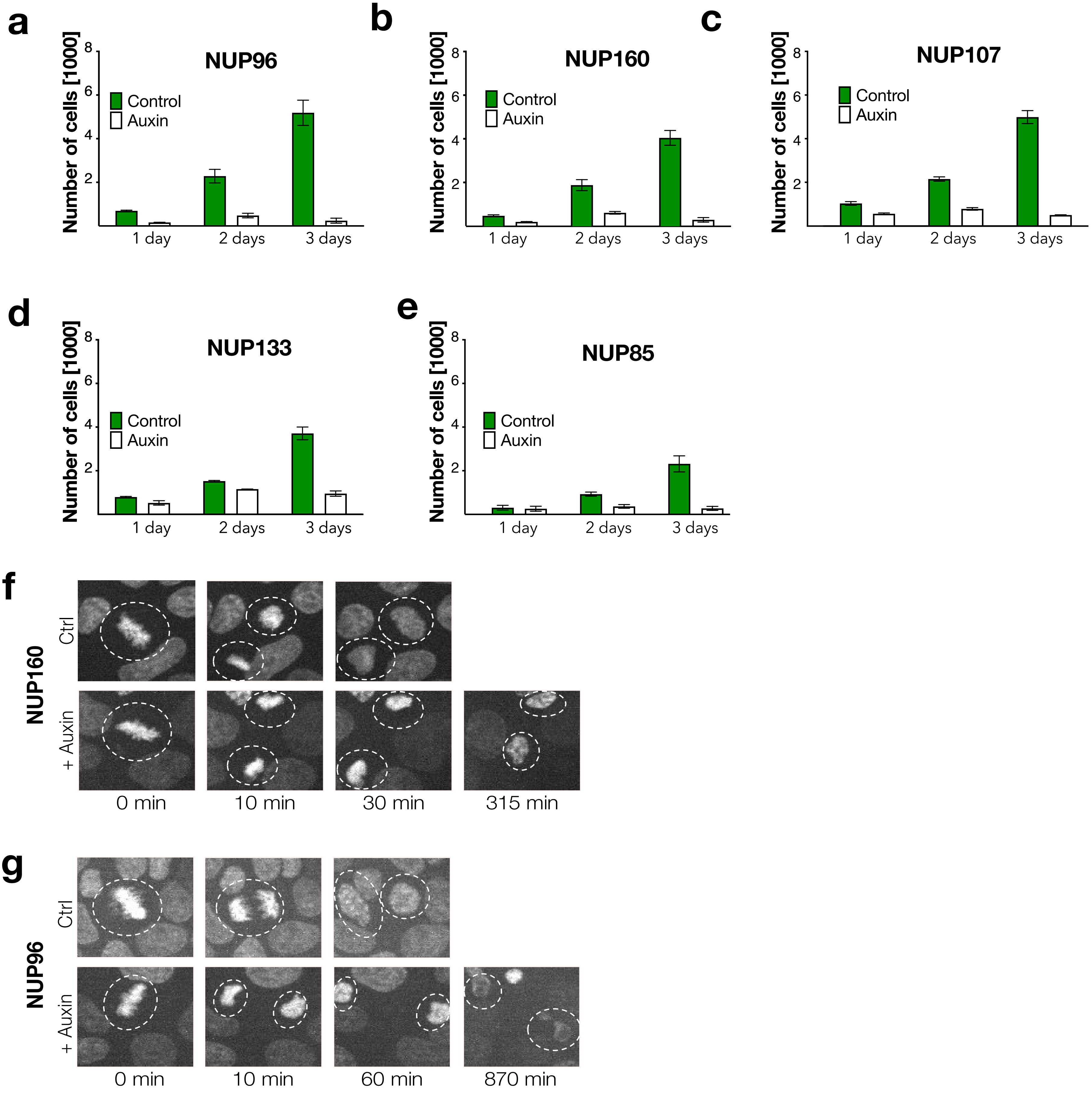

### Supplementary Figure 3. Stability of nuclear pore upon rapid loss of Y-NUPs

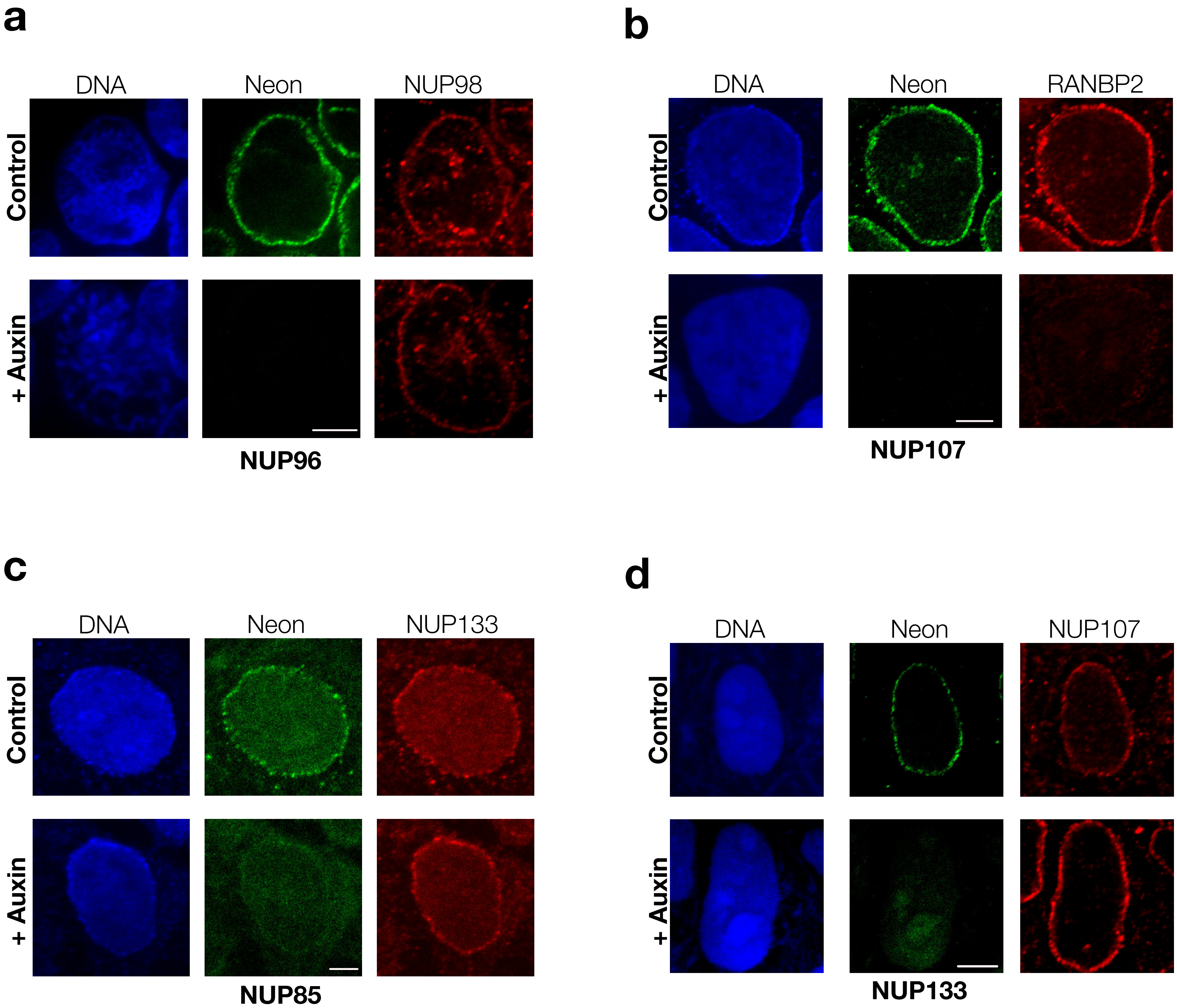

### Supplementary Figure 4. Data supporting NUP93 and NUP188 tagging

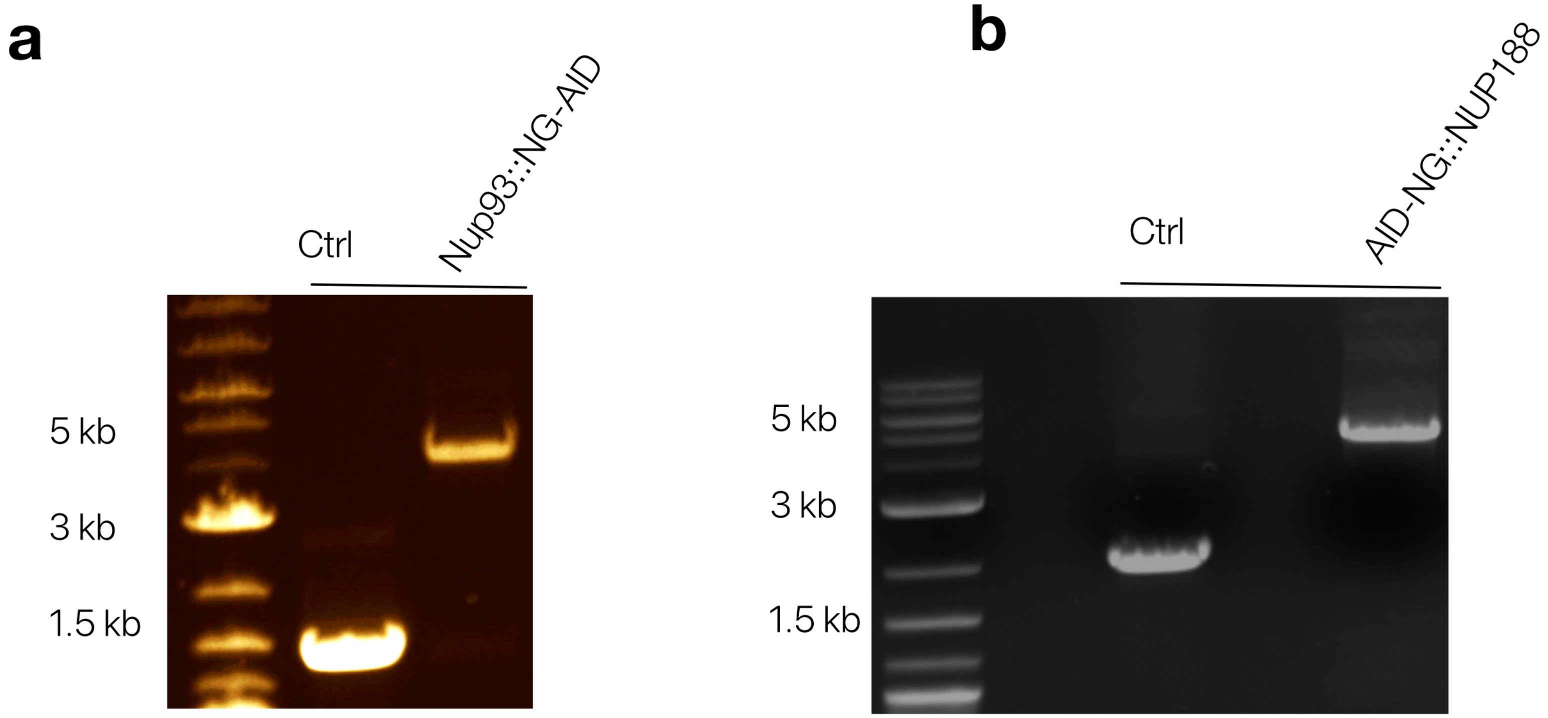

### Supplementary Figure 5. SEM imaging of an individual nucleus

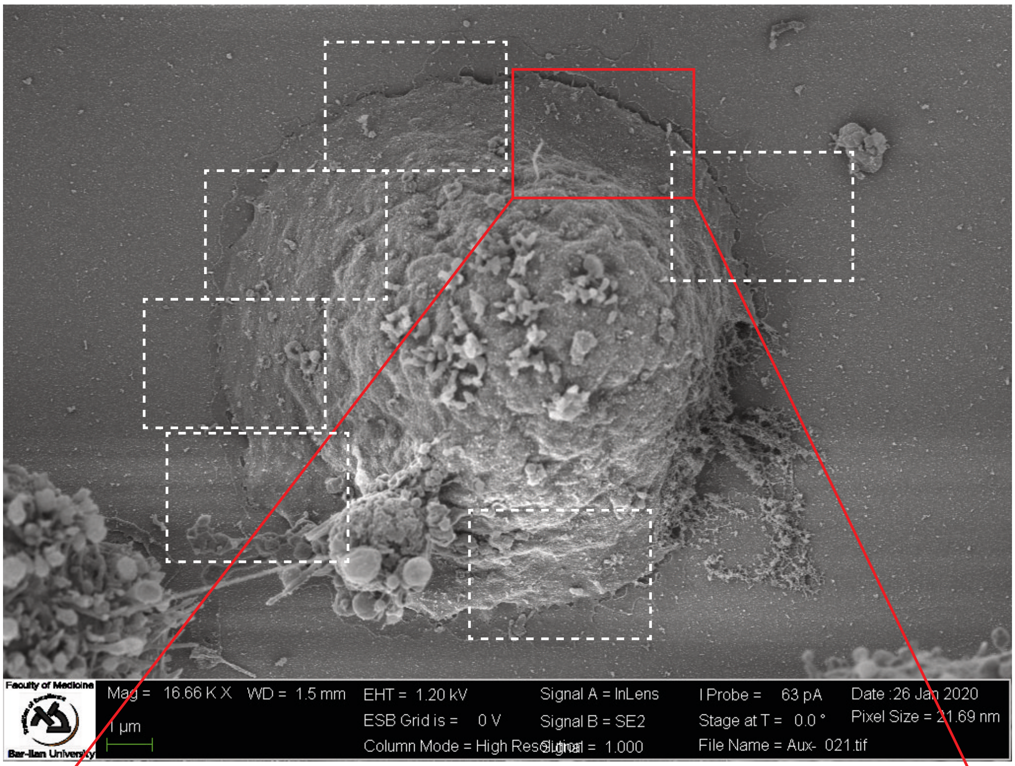

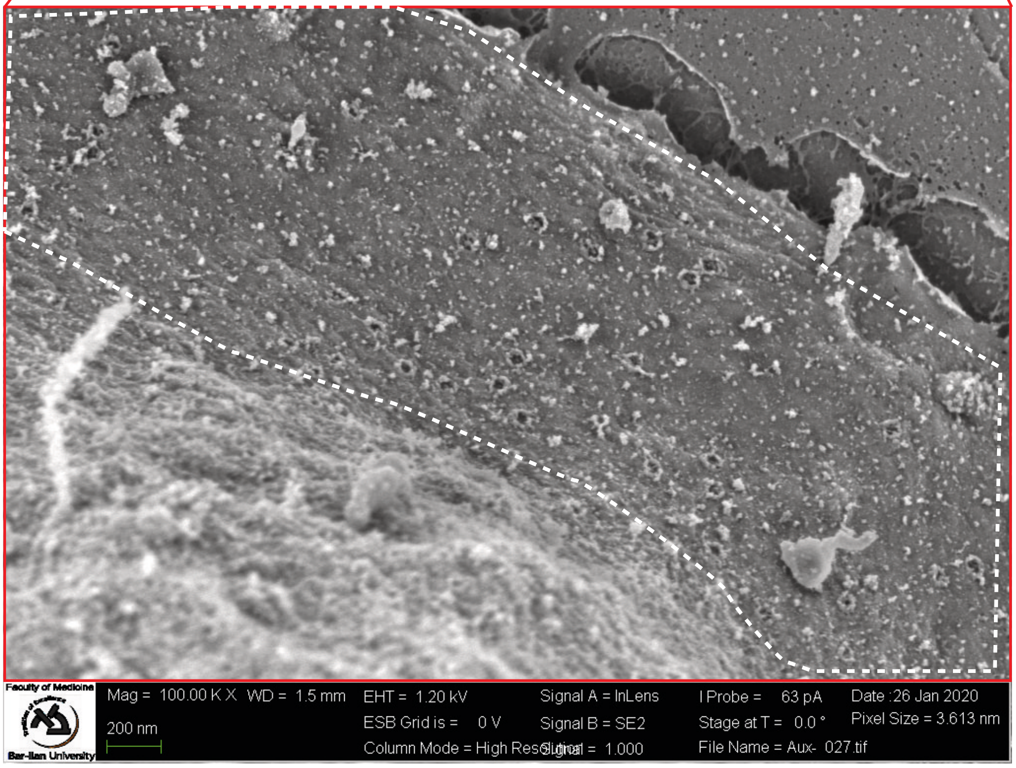

### Supplementary Figure 6. Diversity of structures upon individual nucleoporin depletion

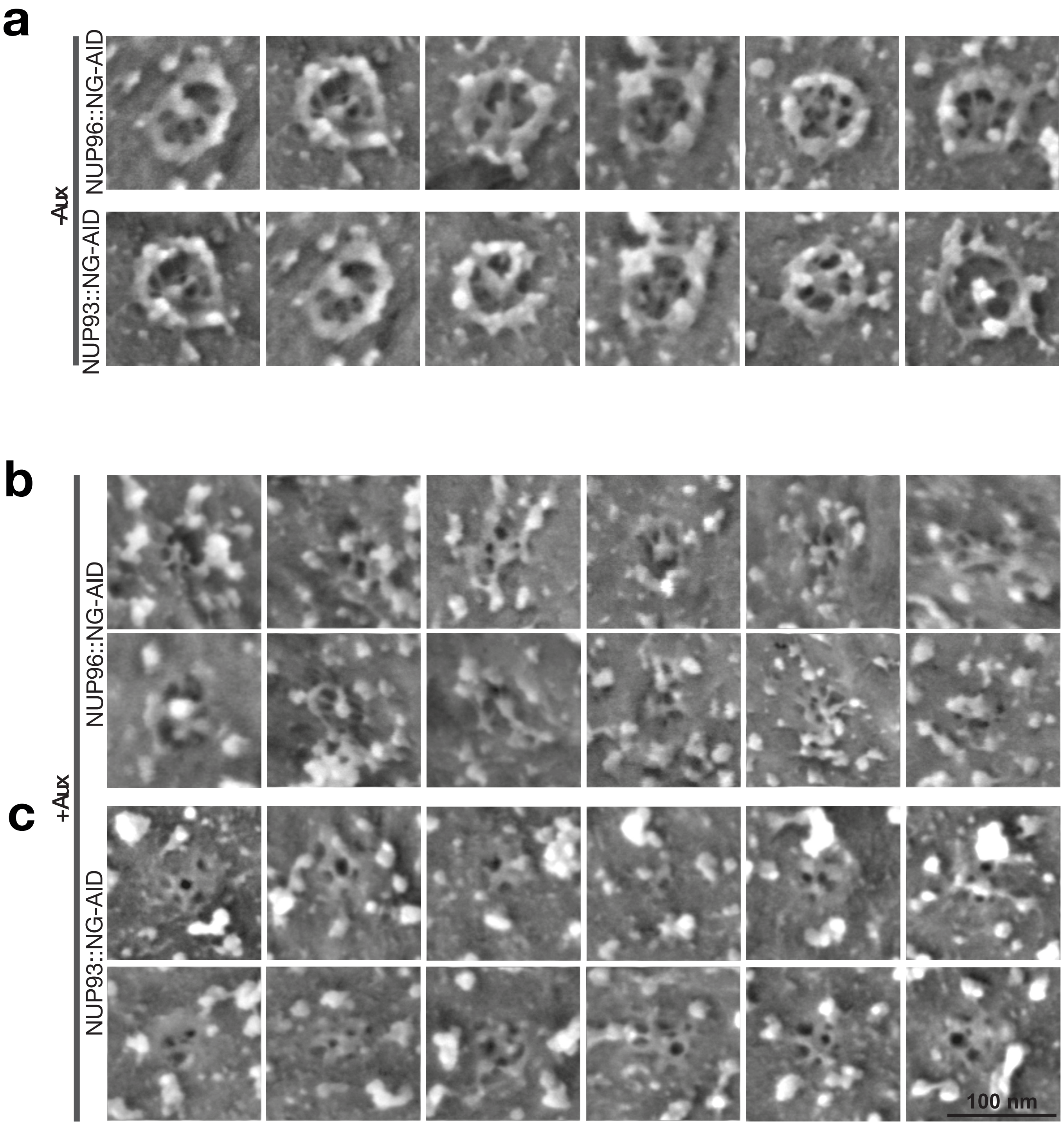
